# A corpus for differential diagnosis: an eye diseases use case

**DOI:** 10.1101/2021.05.10.443343

**Authors:** Antonio Jimeno Yepes, David Martinez Iraola, Pieter Barnard, Tinu Theckel Joy

## Abstract

We have created a corpus for the extraction of information related to diagnosis from scientific literature focused on eye diseases. It was shown that the annotation of entities has a relatively large agreement among annotators, which translates into strong performance of the trained methods, mostly BioBERT. Furthermore it was observed that relation annotation in this domain has challenges, which might require additional exploration.

When using the trained models on MEDLINE, we could identify confirmed knowledge about the diagnosis of eye diseases and relevant new information, which supports the developments in this work. The corpus that we have developed is publicly available, thus the scientific community is able to reproduce our work and reuse the corpus in their work.

## 1. Introduction

The rate at which new information is added to the existing vast amount of scientific literature is ever increasing. Scientific literature databases such as MEDLINE grow by over 1 million new documents every year, which makes it challenging to keep up with the rate of new discoveries and novel methods presented. It is therefore important to focus on methods that can effectively extract and utilise this information. The latter is the theme of this paper, exploring methods for the extraction of entities and their relations from scientific literature.

Our research interests lie in differential diagnosis, in particular differential diagnosis of ocular diseases. That is, the ability to distinguish/differentiate between diseases that may present with similar symptoms. Literature contains many examples of the symptoms associated with different ocular diseases as well as the guidelines and tools used in their diagnosis. We envision an approach where this vast amount of knowledge available in the scientific literature is utilised for medical diagnosis.

Medical guidelines are commonly used to asses diseases and make a classification of a disease degree. Our approach considers all such available guidelines in the scientific literature. More importantly we aim to accelerate the process going from discovering new insights in the domain to their inclusion in medical diagnosis. We therefore present methods related to a data driven approach to medical diagnosis.

According to the World Health Organisation (WHO) ^1^, at least 2.2 billion people have near or distant vision impairment, from which almost half could have been prevented or are not yet addressed. This has a significant impact on the quality of life of individuals suffering from these diseases that could be prevented if diagnosed and treated early on. There is also the economic impact on both governments and private institutions associated with advanced patient care in severe disease cases. Thus, early diagnosis utilising the latest available scientific knowledge is key in early detection and prevention of further disease progression.

Though this article is primarily focused on ocular diseases, recent research has shown promising results in applying the same diagnostic tools for the detection of neurological conditions. Diagnostic tools associated with ocular diseases are of lower cost and thus can be made more readily available than those dedicated to diagnosing neurological conditions. Furthermore, it offers in many cases a non-invasive approach to disease diagnosis. As with ocular diseases, the published scientific literature can provide the needed information for a data driven approach to diagnosing neurological conditions.

In this paper, we present the first manually annotated corpus, that we have made available^2^, for the extraction of diagnostic information from the scientific literature, which has been focused on eye diseases. We evaluate a set of information extraction techniques to reproduce the annotations automatically and we use the trained information extraction methods on a selection of citations from MEDLINE for different eye diseases.

## 2. Related work

The annotation of entities and their relations in corpora is the first approach to develop information extraction methods for specific domains. There are multiple data sets in the biomedical domain that provide this resource, but their focus tends to be on biology or biological processes. Some of these resources have been released in the context of biomedical challenges, such as Biocreative [9], BioNLP [6], and BIOASQ [19]. Another initiative for large-scale annotation (CRAFT) has annotated full documents using different ontologies [1], but the focus is again on biological processes, and not on diseases and their diagnosis.

Previous work by [10, 7] investigated disease entity extraction, though the entities in the current work are studied here for the first time. In recent work, MedMentions [14] has been annotated with entities linked to UMLS (Unified Medical Language System) [4] concepts.

To the best of our knowledge, there are no corpora available relevant to our study that contains manually-annotated relations between the entity types of interest. Examples of relevant previous work include Semantic MEDLINE [11], which contains different types of relations, some of which could be aligned to our study (e.g. “Predisposes”); however after an initial study Semantic MEDLINE did not fit our purposes.

Based on these findings, we have developed a manually annotated data set with entities and relations relevant to diagnosis of eye diseases. In the following section, we describe the entity and relation types defined for our study and how the documents were selected from PubMed and manually annotated.

## 3. Methods

In this section, we describe how we have developed the corpus for diagnosis, starting with the preparation of the annotation guidelines and the manual annotation process. Next, we present several existing methods for named entity recognition and relation extraction, which includes state-of-the-art methods that have been applied on our corpus.

### 3.1. Annotation guidelines

Manual annotations were performed based on a set of predefined annotation guidelines. These guidelines were constructed and subsequently refined based on an initial set of 10 citation documents annotated by three researchers in the domain. These documents were obtained from MEDLINE and were focused primarily on *aged-related macular degeneration* (AMD).

Based on these 10 initial citations, a PubMed template query was created to recover similar MEDLINE citations. This template query contained the name of a disease and at least one of the following *MeSH Qualifiers*: *classification, diagnosis* or *epidemiology*. An example query would be: *“Macular degeneration”[MH] AND (classification[MH] OR diagnosis[MH] OR epidemiology[MH])*. The PubMed searches were constrained to journals with the term Ophthalmology in their name, which includes *Ophthalmology* and *Archives of Ophthalmology*.

Using the template query above we recovered citations for the following diseases that were searched as MeSH Headings: *eye diseases, diabetic retinopathy, glaucoma* and *macular edema*. We used these queries on MEDLINE and randomly selected 50 citations from each set. After removing citations which contained only a title, we selected 195 citations for manual annotation.

Following the alignment and standardisation between annotators as well as defining the needs of differential diagnosis, the following entity and relation types were identified for annotation.

#### 3.1.1. Entity types

**Disease:** a particular abnormal condition that affects part or all of an organism not caused by external force (e.g. injury) and that consists of a disorder of a structure or function, usually serving as an evolutionary disadvantage. Examples of diseases include “AMD” or “glaucoma”.

**Symptom:** a departure from normal function or feeling which is apparent to a patient, reflecting the presence of an unusual state, or of a disease^3^.

**Anatomical part:** Example of anatomical parts include higher level organs, e.g. *eye*, or more fine grained entities such as the different layers that can be observed in the retina. The idea is to identify different areas in which a disease might be present.

**Characteristic:** This entity type annotates characteristics from the population under study that can define the risk factors of a disease.

**Diagnostic tool:** Several tools are used to identify pathologies in patients, e.g. *OCT*, and fundus images. With this entity type, we would like to capture the tools that are used for the identification of pathologies that will lead to diagnosis.

**Dimension:** Some characteristics of the symptoms are defined by measurable properties, such as drusen size that defines from small to large drusens that are measurable from eye images. These dimensions are important since they will allow identifying links between measurable pathologies identified from imaging modalities to the knowledge base. For example a medium drusen dimension is defined as a drusen with a diameter of *>*= *63 and < 125 μm*.

**Classification System:** Diagnosis is typically done using existing guidelines or international recommendations. This entity type intends to capture the classification systems identified in the papers.

#### 3.1.2. Relation types

**Alias:** Relates entities that have the same meaning but different surface form (e.g. p21WAF1 and p21 in MEDLINE abstract with PubMed id 25275039).

**Is a:** This relation denotes that an entity is more specific in comparison to another entity in the document. As with the Alias relation, annotations should be done to the first occurrence of the relation in an abstract and it can span across sentences.

**Causes:** Relates diseases and symptoms or diseases when it is known in the context what disease causes a symptom or disease.

**Has symptom:** Diseases might have presentations that can be linked to a specific symptom.

**Has risk factor:** Relate a disease to a characteristic.

**Has dimension:** Relates a symptom and its dimension, e.g. drusen size.

**Diagnosed with:** Tool or classification system used to diagnose or grade a disease.

**Located in:** Relates a disease and the anatomical location it might be in.

### 3.2. Manual corpus annotation

Annotations were performed by four annotators using the BRAT tool [18]. Each citation was annotated by two annotators and then merged automatically. This merging process would show where the disagreement between annotators was if any. The disagreement between annotators was resolved after the merge. A total of 195 citations were annotated, which includes the calibration documents.

We calculated the inter-annotator agreement using the doubly annotated documents. Precision, recall and F1 [20] were calculated as shown in Table 1. An exact match of the entity boundaries (Exact) and a more relaxed version (Partial) have been estimated using BRATEval^4^ [20].

**Table 1:**
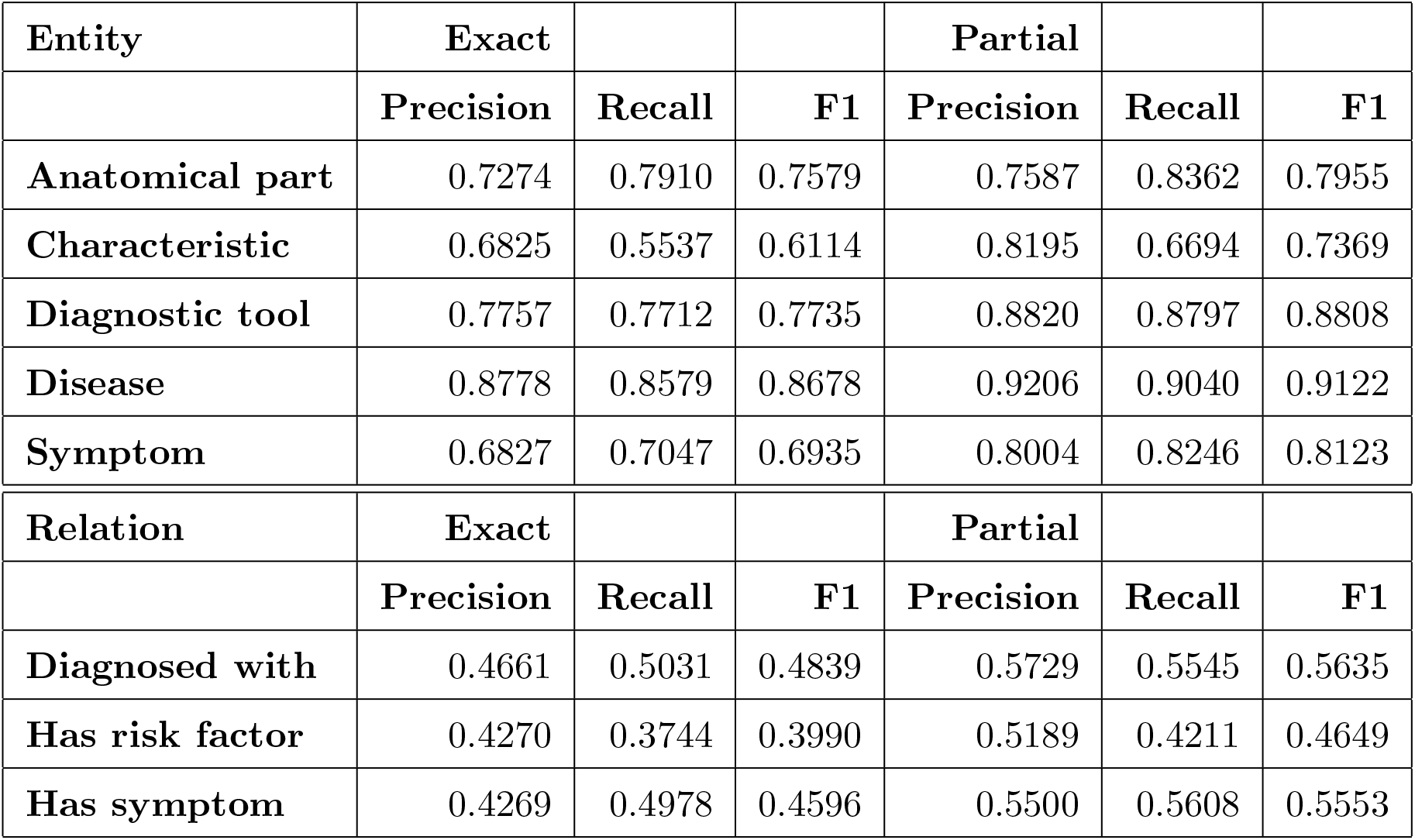
Inter-Annotator Agreement from our manually annotated set

| **Entity** | **Exact** |  |  | **Partial** |  |  |
| --- | --- | --- | --- | --- | --- | --- |
|  | **Precision** | **Recall** | **F1** | **Precision** | **Recall** | **F1** |
| **Anatomical part** | 0.7274 | 0.7910 | 0.7579 | 0.7587 | 0.8362 | 0.7955 |
| **Characteristic** | 0.6825 | 0.5537 | 0.6114 | 0.8195 | 0.6694 | 0.7369 |
| **Diagnostic tool** | 0.7757 | 0.7712 | 0.7735 | 0.8820 | 0.8797 | 0.8808 |
| **Disease** | 0.8778 | 0.8579 | 0.8678 | 0.9206 | 0.9040 | 0.9122 |
| **Symptom** | 0.6827 | 0.7047 | 0.6935 | 0.8004 | 0.8246 | 0.8123 |
| **Relation** | **Exact** |  |  | **Partial** |  |  |
|  | **Precision** | **Recall** | **F1** | **Precision** | **Recall** | **F1** |
| **Diagnosed with** | 0.4661 | 0.5031 | 0.4839 | 0.5729 | 0.5545 | 0.5635 |
| **Has risk factor** | 0.4270 | 0.3744 | 0.3990 | 0.5189 | 0.4211 | 0.4649 |
| **Has symptom** | 0.4269 | 0.4978 | 0.4596 | 0.5500 | 0.5608 | 0.5553 |

**Table 2:**
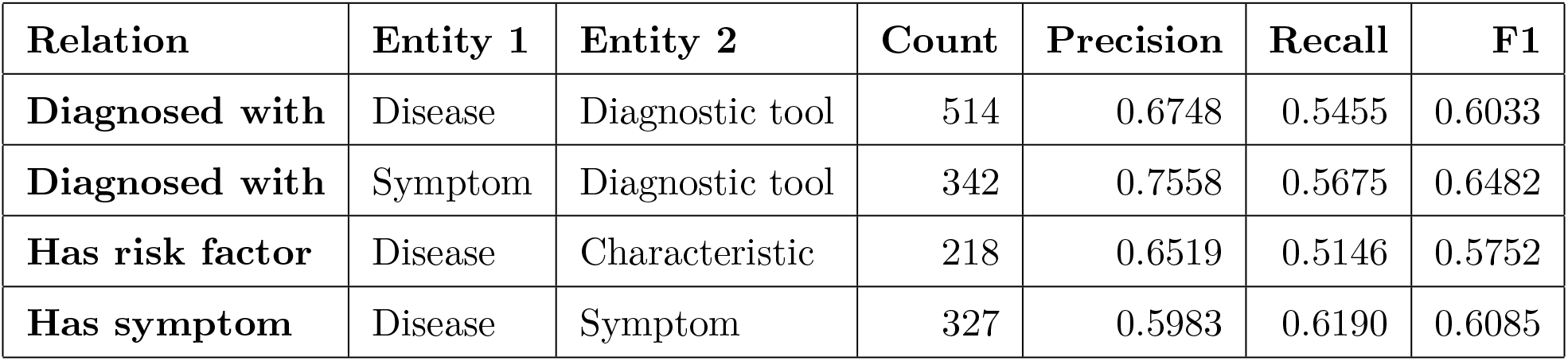
Inter-annotator agreement of manual relations when entities agree

| **Relation** | **Entity 1** | **Entity 2** | **Count** | **Precision** | **Recall** | **F1** |
| --- | --- | --- | --- | --- | --- | --- |
| **Diagnosed with** | Disease | Diagnostic tool | 514 | 0.6748 | 0.5455 | 0.6033 |
| **Diagnosed with** | Symptom | Diagnostic tool | 342 | 0.7558 | 0.5675 | 0.6482 |
| **Has risk factor** | Disease | Characteristic | 218 | 0.6519 | 0.5146 | 0.5752 |
| **Has symptom** | Disease | Symptom | 327 | 0.5983 | 0.6190 | 0.6085 |

The final corpus contains 195 annotated citations. There are 1753 mentions of *diseases*, 1501 mentions of *characteristics*, 1493 mentions of *diagnostic tools*, 1257 mentions of *symptoms*, 907 mentions of *anatomy parts*, 95 mentions of *classification systems* and 34 mentions of *dimensions*. Table 3 shows the 10 most frequent terms per entity type, which seems to align with what would be expected in the domain of the retrieved MEDLINE citations.

**Table 3:**
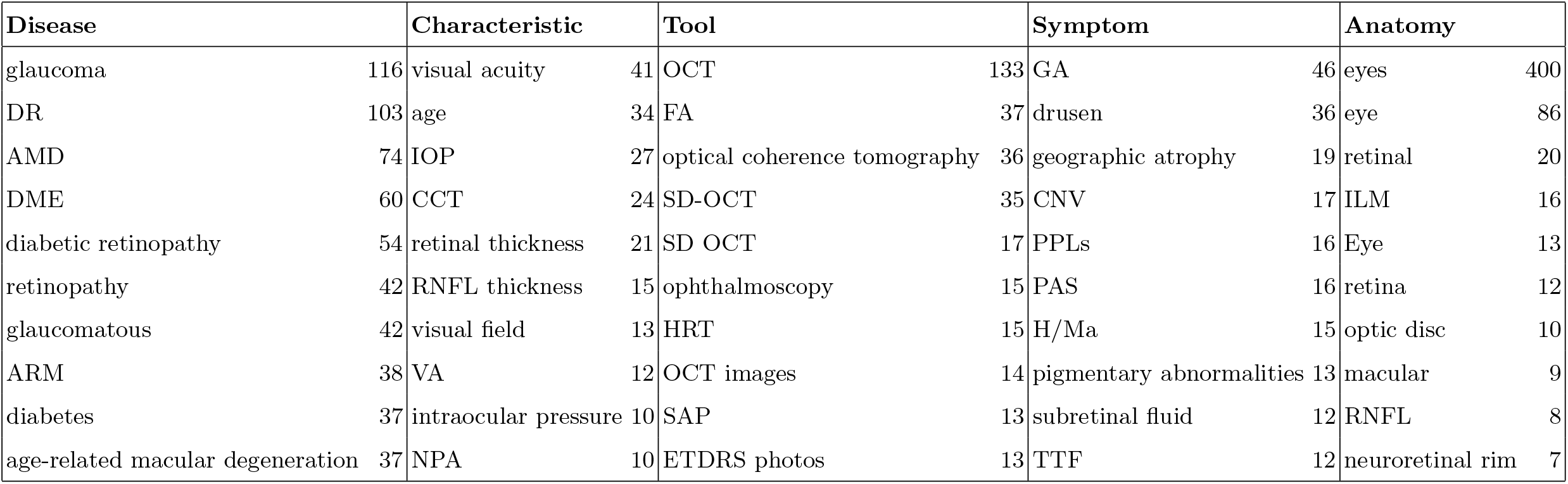
Top 10 most frequent terms per entity type in our manually annotated corpus

There are 931 mentions of *diagnosed with* relations, 415 *has symptom* relations, 276 *has risk factor* relations, 94 *causes*, 61 *associated with*, 19 *located in* and 18 *has dimension*. There as well negative relations, even though their number is less frequent compared to positive ones. We found 15 *no has risk factor* relations, 11 *not associated with* relations, 11 *no has symptom* relation, 5 *no diagnosed with* and 5 *no causes*. We identify 505 *alias* relations and 412 *is a* relations, which will support learning information about the language used in these documents and taxonomic relations.

### 3.3. Named entity recognition

We have evaluated two machine learning methods for named entity recognition. The first method is the Conditional Random Field (CRF) model using engineered features, while the second method is the Bidirectional Encoder Representations from Transformers (BERT) model trained on biomedical data, namely the BioBERT model.

#### 3.3.1. Conditional Random Field

Similar to previous work [10], we have evaluated a Conditional Random Field based annotator on our data set. Our implementation relies on UIMA[8] (https://uima.apache.org) with uimaFit (https://uima.apache.org/uimafit.html) as the Natural Language Processing (NLP) framework and ClearTK[2] (https://cleartk.github.io/cleartk) as the UIMA machine learning package and CRFSuite[16] as the CRF implementation, for which an UIMA wrapper was developed. No specific hyper-parameter tuning has been made.

Text is split into sentences using ClearTK. For training, the text is tokenized and the tokens are assigned a BIO (Beginning/In/Out of entity) label. Each token is represented using its surface form, 2/3-gram characters from the end of the token, a lower cased form and two features indicating if the term is in capital letters and if it is a number, which are relevant in the biomedical domain.

#### 3.3.2. BioBert NER

BioBERT^5^ [13] is a pre-trained BERT model trained on biomedical data. More specifically we have used the BioBERT model pre-trained on both the PubMed as well as PubMed central datasets, namely the *biobert_v1*.*0_pubmed model*. BioBERT has a transformer encoder network with a fully connected layer added for classification of entity tokens. In our case the classification predicts the BIO (Beginning/In/Out of entity) encoding of the entities. The BIO output is then converted back to the BRAT annotation format to allow for all methods to be evaluated using the same format. The standard BioBERT code was modified to allow for the annotation of multiple entity types.

### 3.4. Relation extraction

For relation extraction, we have considered three machine learning methods, one based on a Support Vector Machine (SVM) classifier with engineered features and two other, both based on state-of-the-art deep learning approaches, namely *TEES (Turku Event Extraction System)* and *BioBERT*.

#### 3.4.1. SVM with engineered features

In this method, we have engineered a set of features that we have used to train a SVM to predict if two entities are related in a sentence. We use LibSVM[5] with a linear kernel as the SVM implementation and the same underlying NLP framework as the CRF method for named entity recognition.

For training, we have built sets of positive and negative instances. Positive instances are identified by entities that appear as related in the ground truth. Negative instances are identified by entities of the types of the relation arguments that are not related.

Each instance is represented as a set of clearly distinguished features that include the distance in tokens between the entities, the tokens in the entities, and the tokens between the entities. We investigated more complex features derived from syntactic parsing, but there were no benefits derived from them.

#### 3.4.2. TEES

Turku Event Extraction System (TEES) is a free and open source NLP system developed for the extraction of entities, relations, and events from biomedical text [3]. For this work we rely on the relation classification step of the pipeline. The input examples are modelled in the context of a sentence window, centered around the candidate relation, and the sentence is modelled as a linear sequence of word tokens. Each token is mapped to different vector space embeddings, based on: word vectors, part-of-speech, entity, distance, relative position, and dependency paths. These embeddings are concatenated together, resulting in an n-dimensional vector, which is processed by a set of 1D convolutions with window sizes 1, 3, 5 and 7. Global max pooling is applied for each convolutional layer and the resulting features are merged together into the convolution output vector.

This output vector is fed into a dense layer of 200 to 800 neurons, which is connected to the final classification layer where each label is represented by one neuron. The classification layer uses sigmoid activation, and the other layers use ReLU [15] activation. Classification is performed as multi-label classification where each example may have 0 to n positive labels. They use the Adam optimizer [12] with binary cross-entropy and a learning rate of 0.001.

In addition to TEES, we have used a modified version [17] based on multi-head attention. The base implementation uses TEES but the CNN implementation is modified by multi-head attention with 4 heads. This modification intends to deal with long dependencies, which might be missed by the CNN method as shown in [17].

#### 3.4.3. BioBERT

BioBERT [13] is a pre-trained language model that was used to model the relation between two given entities. Fine-tuning was performed by allowing the pre-trained model to be tuned on a data set specific to the domain of interest. Entities in this data set were masked in the input sentences by tagging it with a preceding @ and an ending $, e.g. @Disease$. Thus, the model aims to gain context of the relation between masked entities by using non-masked words in the text. The token representation in the encoder output layer is then subsequently fed to a linear layer for classification of the relation. A model was built for each of the relations given in Table 1 and thus binary classification was performed to determine the presence or absence of the given relation.

## 4. Results

In this section, we present the result for the automatic reproduction of the manual annotations. We show results for entity recognition first, followed by the relation extraction experiments.

### 4.1. Named entity recognition (NER)

Table 4 shows the results of the CRF and BioBERT methods. The performance of the BioBERT model is higher than the CRF method. Interestingly, except for diseases and anatomical parts, there is a large increase in performance between the exact and partial results. In the case of BioBERT, the performance is very similar to the inter-annotator performance, which could be considered as the upper bound performance of NER methods on this data set.

**Table 4:** Automatic entity annotation results

| **CRF** | **Exact** |  |  | **Partial** |  |  |
| --- | --- | --- | --- | --- | --- | --- |
|  | **Precision** | **Recall** | **F1** | **Precision** | **Recall** | **F1** |
| **Anatomical part** | 0.8762 | 0.4071 | 0.5559 | 0.9048 | 0.4213 | 0.5749 |
| **Characteristic** | 0.2623 | 0.1427 | 0.1848 | 0.6189 | 0.4092 | 0.4927 |
| **Diagnostic tool** | 0.5389 | 0.3165 | 0.3988 | 0.8741 | 0.5918 | 0.7058 |
| **Disease** | 0.8262 | 0.5827 | 0.6834 | 0.9238 | 0.6649 | 0.7732 |
| **Symptom** | 0.2916 | 0.1881 | 0.2287 | 0.6758 | 0.5258 | 0.5915 |
| **BioBERT** | **Exact** |  |  | **Partial** |  |  |
|  | **Precision** | **Recall** | **F1** | **Precision** | **Recall** | **F1** |
| **Anatomical part** | 0.6972 | 0.7821 | 0.7372 | 0.7525 | 0.8366 | 0.7923 |
| **Characteristic** | 0.5618 | 0.6247 | 0.5915 | 0.7086 | 0.7700 | 0.7381 |
| **Diagnostic tool** | 0.6932 | 0.7560 | 0.7232 | 0.8312 | 0.8904 | 0.8598 |
| **Disease** | 0.8523 | 0.8775 | 0.8647 | 0.9040 | 0.9269 | 0.9153 |
| **Symptom** | 0.6172 | 0.6949 | 0.6538 | 0.7749 | 0.8452 | 0.8085 |

### 4.2. Relation extraction

Table 5 shows the results for the relation extraction methods^6^. We can see that for all relation types, except for *Has risk factor*, the performance is better than the inter-annotator agreement (cf. Table 2), this is because we provided the entities and the task consists of only predicting the relations. The best scores are achieved for the *Diagnosed with* relation (the ones with most training data). For the *Has risk factor* relation all classifiers perform poorly with F1 scores below 0.40.

**Table 5:** Automatic relation extraction results

| **SVM** | **Entity 1** | **Entity 2** | **Precision** | **Recall** | **F1** |
| --- | --- | --- | --- | --- | --- |
| **Diagnosed with** | Disease | Diagnostic tool | 0.5948 | 0.8220 | 0.6902 |
| **Diagnosed with** | Symptom | Diagnostic tool | 0.6575 | 0.8638 | 0.7467 |
| **Has risk factor** | Disease | Characteristic | 0.3917 | 0.3696 | 0.3803 |
| **Has symptom** | Disease | Symptom | 0.5647 | 0.7654 | 0.6499 |
| **TEES** | **Entity 1** | **Entity 2** | **Precision** | **Recall** | **F1** |
| **Diagnosed with** | Disease | Diagnostic tool | 0.6142 | 0.8924 | 0.7276 |
| **Diagnosed with** | Symptom | Diagnostic tool | 0.6780 | 0.8399 | 0.7503 |
| **Has risk factor** | Disease | Characteristic | 0.4318 | 0.3180 | 0.3663 |
| **Has symptom** | Disease | Symptom | 0.5516 | 0.7051 | 0.6190 |
| **TEES Multi-head** | **Entity 1** | **Entity 2** | **Precision** | **Recall** | **F1** |
| **Diagnosed with** | Disease | Diagnostic tool | 0.6071 | 0.8767 | 0.7174 |
| **Diagnosed with** | Symptom | Diagnostic tool | 0.6882 | 0.7822 | 0.7322 |
| **Has risk factor** | Disease | Characteristic | 0.3744 | 0.3054 | 0.3364 |
| **Has symptom** | Disease | Symptom | 0.5757 | 0.8118 | 0.6737 |
| **BioBERT** | **Entity 1** | **Entity 2** | **Precision** | **Recall** | **F1** |
| **Diagnosed with** | Disease | Diagnostic tool | 0.6797 | 0.7177 | 0.6982 |
| **Diagnosed with** | Symptom | Diagnostic tool | 0.7136 | 0.7223 | 0.7179 |
| **Has risk factor** | Disease | Characteristic | 0.4651 | 0.2578 | 0.3317 |
| **Has symptom** | Disease | Symptom | 0.6251 | 0.6337 | 0.6294 |

## 5. Discussion

In this section, we are going to analyse the results for entity recognition and relation extraction and will apply the trained models on a set of diseases to understand the findings based on reusing these models.

### 5.1. Automatic annotation

#### Entity extraction

Comparing a Conditional Random Field (CRF) model and a BERT based one (BioBERT), we find that the BERT based model has higher F1 scores and can thus be said to perform better. The BioBERT model results show a similar trend as observed for the inter-annotator results in Table 1, with the highest F1 score obtained for the *Disease* entity and the lowest F1 score for the *Characteristic* entity.

For both models, the *Characteristic* category is the most challenging one. We manually analysed the results of a sample of predictions for this entity in order to determine the main reasons for the low performance. With regards to False Positives (FPs), we found that the most common source of error (31 out of 82 FPs) is the annotation of general terms that are not well specified characteristics of a population (e.g. “morphologic characteristics”, “family history”, etc.). Closely following, another main source of FPs (24/82) was caused by the mismatch with the boundaries of the entity. The 3rd main cause of FPs was the confusion with the entity *Symptom* (7/82).

We also manually checked False Negatives (FNs), and found that the main source of error was boundary mismatch (20/45), followed closely by confusion with other entities (19/45). In particular the entity *Symptom* led to 12 FNs, and this illustrates the high confusability between these entity types.

#### Relation extraction

The respective performance metrics in Table 5 indicate similar performance for the different models. The scores are in line with what was observed for the inter annotator results, with low F1 values for the *Has risk factor* relation. Relation extraction outcomes are not as high as with entity recognition, but follow a similar performance to previous work [20].

In general, there is little difference between the respective performance metrics between the models. This is surprising to some extent as BERT based models are expected to outperform classical ML methods like SVM. However, considering the size of the data set this could be explained. We also observe that BioBERT has higher precision values than other classifiers, although at the cost of lower recall. Looking at the TEES based methods, the *Has symptom* relation for the multi-headed attention approach shows a clear improvement over the convolution based equivalent.

In order to better understand the low performance, we manually analysed a sample of results for the relation types *Has risk factor* and *Has symptom*, consisting of all errors and true positives from one of the folds for the classifier TEES. We start with *Has risk factor*, and present some examples to discuss in Figure 2. With regards to false positives, we noticed that a large proportion (16/29) correspond to cases where there is no explicit mention that the population is a risk factor, but it is under study. For instance, in sentence (a) in Figure 2, we find that the classifier has mistakenly predicted the unproven relation between the population group and the disease under study. Another repeated false positive is the case of negated relations (3/29), which do not have enough data to be trained separately, and are not processed in any other way (cf. example (b) in Figure 2).

**Figure 1:**
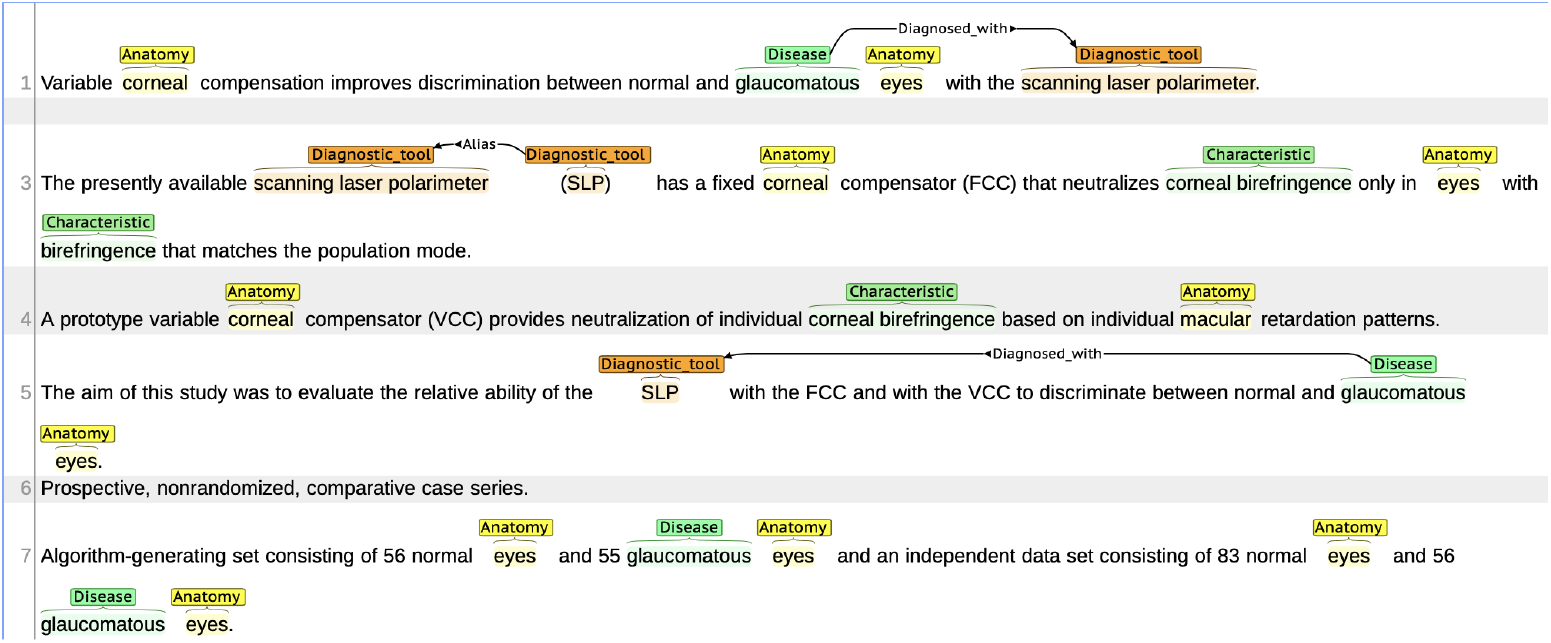
Example annotation screenshot of citation with PMID:15019373

**Figure 2:**
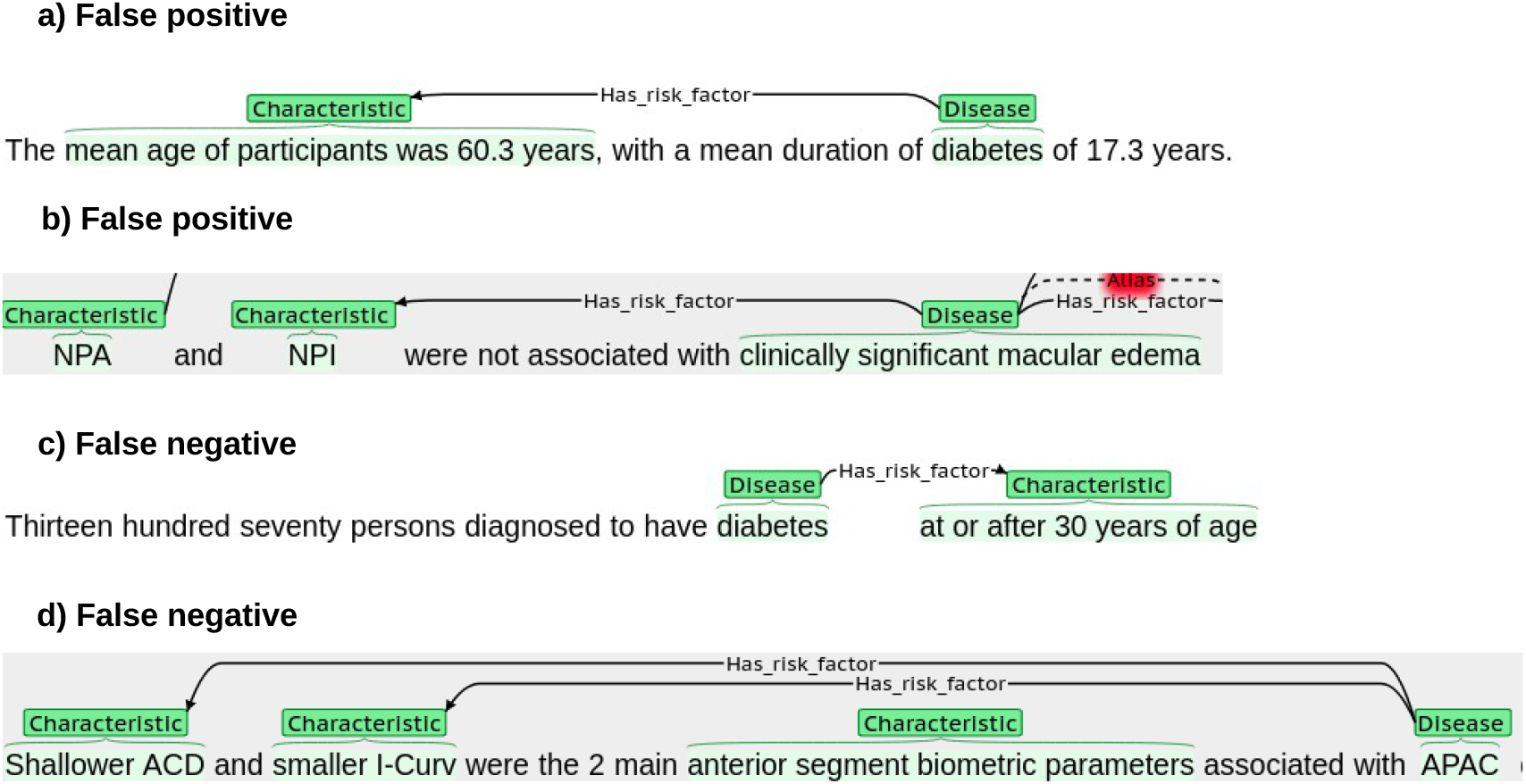
Examples of false positives and false negatives for relation type *Has risk factor* and classifier TEES. False positives show predictions by TEES, and false negatives show gold annotation missed by TEES.

We moved on to verify false negatives, and found that in 3 out of 11 cases, they correspond to population groups that are being studied, without explicit mention that a *Has risk factor* relation exists (cf. example (c) in Figure 2). The confusion in the annotation of these cases could explain the low inter-annotator agreement and the low performance of the different classifiers, suggesting that the annotation of this category should be revisited. The remaining false negatives do not seem to show a clear pattern, although in all cases (8/11) the entity *Characteristic* is a kind of body measurement, such as in example (d) in Figure 2. In this example both relations are missed by TEES, although the multi-head attention version is able to predict one of them.

We also analysed relation type *Has symptom*, which presents a much better performance for all classifiers. In this case, for false positives, we observe that a large proportion of these (23/38) are caused by sentences that list symptoms and diseases without explicit relations between them. We can see an example of the manual annotation at the top of Figure 3, where no relation has been identified; however TEES finds multiple false positives linking the symptoms and diseases. With regards to false negatives, we found that in most cases (33/39), the errors appear in sentences with multiple mentions of symptoms and diseases. For instance at the bottom of Figure 3, we can see the predictions of TEES for a sentence with two mentions of diseases, and four of symptoms. TEES correctly identifies the four relations with the first mention, but fails to predict the relations to the second mention, which are captured in the manual annotation.

**Figure 3:**
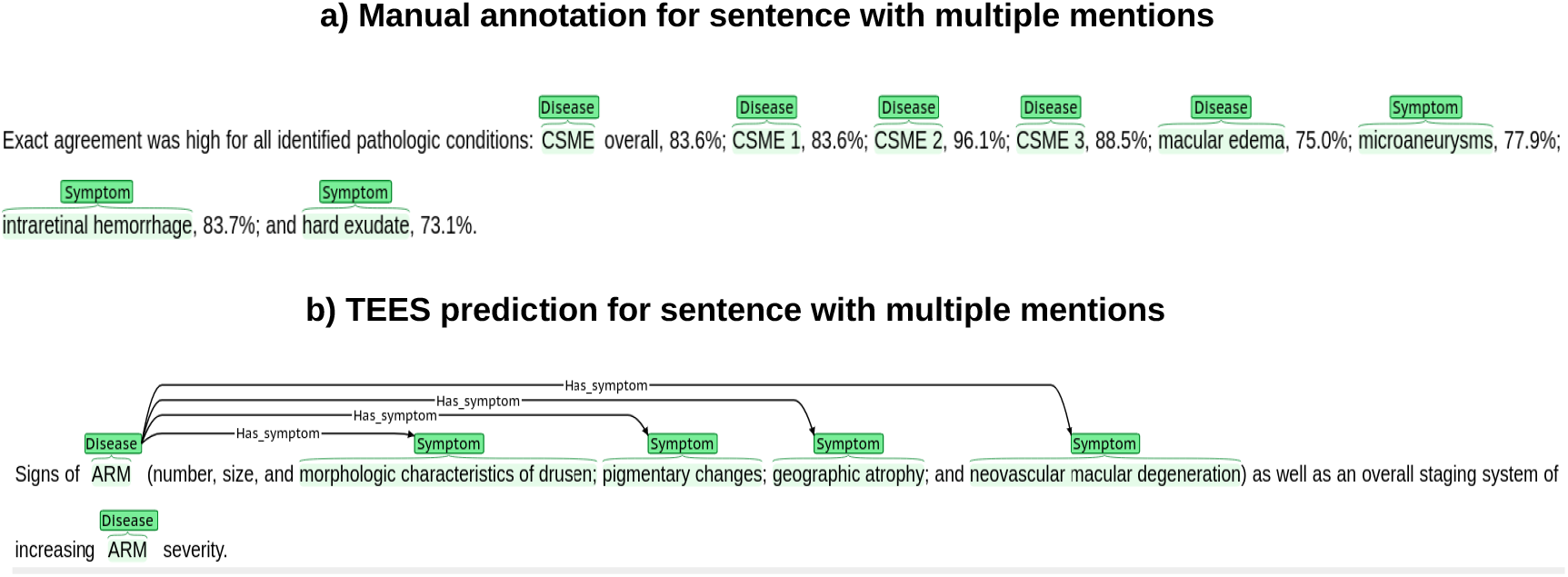
Examples of manual annotation and TEES predictions for example sentences involving relation type *Has symptom*.

The results from the analysis of *Has symptom* suggest that the errors are due to the difficulty of some sentences (with multiple mentions and long dependencies), and not to annotation inconsistencies.

### 5.2. MEDLINE annotation analysis

In addition to automatic annotation, we have done a preliminary evaluation of the BioBERT model for named entity recognition and the SVM models for relation extraction using subsets from MEDLINE. We have collected MED-LINE abstracts for 4 disease groups using the query in the Methods section and aggregated information using the information extraction methods. The 4 disease groups include: eye diseases (as MeSH Heading to select all eye diseases), diabetic retinopathy, glaucoma and macular edema.

Since the names of diseases and other entities might have lexical variants in text, we have used the UMLS 2020AB with level 0 vocabularies to normalise them, e.g. *DME* and *diabetic macular edema* are represented as the same entity. We have used the UMLS Semantic Types to identify concepts in the UMLS that map to the entity types in our work, more precisely to characteristics, diagnostic tools, diseases and symptoms. Entities that could not be mapped to UMLS concepts were not normalized but they are still shown in the analysis.

Results are available in tables 6 to 9 for the *eye diseases* corpus, additional results for the other subsets are available as Supplementary Material. Each table shows results per relation type and the more frequent relations for each disease.

**Table 6:** Relation: Diagnosed with / Disease - Diagnostic tool

| Disease | Diagnostic tool | Frequency |
| --- | --- | --- |
| glaucomas | tomogr optical coherence | 51 |
| diabetic macular edema | tomogr optical coherence | 49 |
| glaucomatous | tomogr optical coherence | 19 |
| macular retinal edema | tomogr optical coherence | 18 |
| glaucomas | visual field study | 12 |
| glaucomas | mtcp1 | 11 |
| glycogen storage disease type ii | tomogr optical coherence | 11 |
| age related macular degeneration | tomogr optical coherence | 10 |
| myopias | tomogr optical coherence | 10 |
| cme | tomogr optical coherence | 10 |

A first look at the most frequent entities shows the most studied diseases and related entities more frequent in our MEDLINE subsets. Among the most common diseases, *glaucoma* and *macular* related diseases seem to be more frequent. Within the symptoms, we identify *choroidal neovascularisation (CNV), geographic atrophy* and *visual defects*, among others. Characteristics includes a diverse set of terms to identify age groups and several measurements (e.g., IOP, intraocular pressure).

As well, we find a confirmation of established knowledge, which is expected. The most common diagnostic tools such as *tomographic optical coherence* and more specific versions of it are very frequent in tables 6 and 7. We identify as well some less common methods for eye disease diagnosis, which includes the gene *MTCP1*, associated with glaucoma. In some cases, therapeutic procedures have been identified as diagnostic tools, such as *hormone replacement therapies* for glaucoma. In tables 8 and 9, we identify similar findings for symptoms and characteristics related the diseases that we are considering in our work.

**Table 7:** Relation: Diagnosed with / Symptom - Diagnostic tool

| Symptom | Diagnostic tool | Frequency |
| --- | --- | --- |
| cnv | doxorubicin/fluorouracil | 22 |
| cnv | octa | 13 |
| hole, macular | tomogr optical coherence | 11 |
| visual field defect | mtcp1 | 10 |
| calcinoses | cat scans, x ray | 10 |
| vitreomacular traction | tomogr optical coherence | 9 |
| field loss | manual perimetry | 8 |
| atrophies | tomogr optical coherence | 8 |
| epiretinal membranes | tomogr optical coherence | 8 |
| atrophy, geographic | sd oct | 7 |

**Table 8:** Relation: Has risk factor / Disease - Characteristic

| Disease | Characteristic | Frequency |
| --- | --- | --- |
| amblyopia | children | 9 |
| amblyopia | preverbal children | 5 |
| strabismus | children | 5 |
| fibroplasias, retrolental | infants | 5 |
| retinoblastomas | children | 5 |
| glaucoma, chronic simple | vf | 4 |
| glaucomas | cone density | 4 |
| amblyopia | preschool children | 4 |
| retinopathy of prematurity, stage 3 | infants | 4 |
| fibroplasias, retrolental | premature infants | 3 |

**Table 9:**
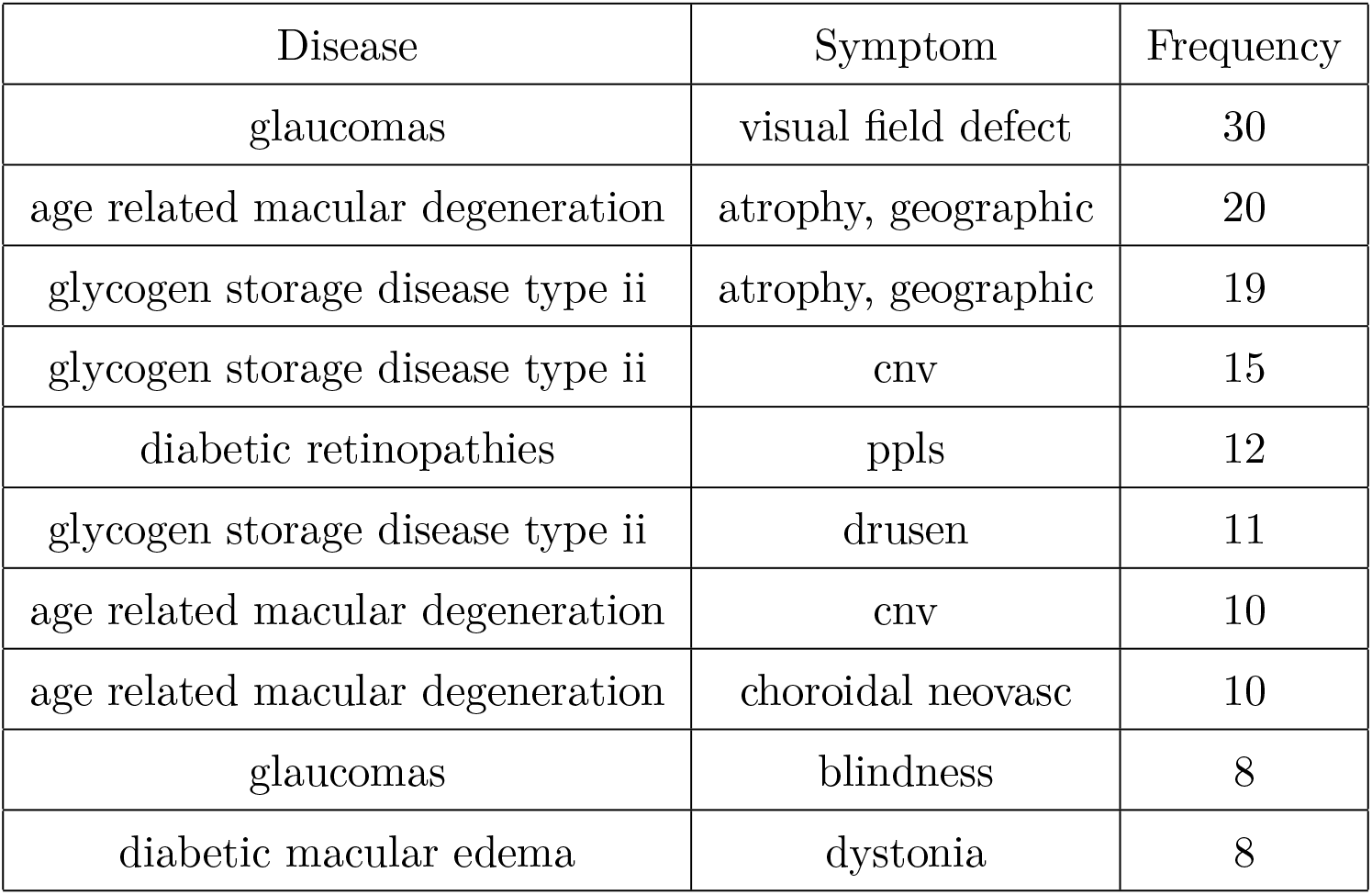
Relation: Has symptom / Disease - Symptom

| Disease | Symptom | Frequency |
| --- | --- | --- |
| glaucomas | visual field defect | 30 |
| age related macular degeneration | atrophy, geographic | 20 |
| glycogen storage disease type ii | atrophy, geographic | 19 |
| glycogen storage disease type ii | cnv | 15 |
| diabetic retinopathies | ppls | 12 |
| glycogen storage disease type ii | drusen | 11 |
| age related macular degeneration | cnv | 10 |
| age related macular degeneration | choroidal neovasc | 10 |
| glaucomas | blindness | 8 |
| diabetic macular edema | dystonia | 8 |

## 6. Conclusions and future work

In this work, we have developed a corpus for the extraction of information related to diagnosis, which we have focused on eye diseases. We have identified that annotation of entities has a relatively large agreement among annotators, which translates in a strong performance of the trained methods, mostly BERT. We observe as well that relation annotation in this domain has challenges, which might require additional exploration. On the other hand, we could identify confirmed knowledge and relevant new information using trained information extraction methods on this corpus, which supports the developments in this work. The corpus that we have developed is publicly available, thus the scientific community is able to reproduce our work and reuse the corpus in their work.

As an extension of our work, we would like to use the trained models in domains other than eye diseases and identify how reusable our effort is. We have done extraction based on citations and it would be worth extending the study to full text articles. Finally, we would like to analyse the new information extracted by the information extraction methods to further understand their significance in the eye diseases domain.

## 7. Acknowledgements

We would like to thank Rahil Garnavi, Bhavna Antony and Yulia Otmakhova for fruitful discussions and suggestions. We would like to thank Alexa Moses for her support and help in releasing the data set.

## Footnotes

1 https://www.who.int/news-room/fact-sheets/detail/blindness-and-visual-impairment

2 Diagnosis Corpus repository: https://github.com/ibm-aur-nlp/diagnosis-corpus

3 https://en.wikipedia.org/wiki/Signs_and_symptoms

4 https://bitbucket.org/nictabiomed/brateval

5 https://github.com/dmis-lab/biobert

6 In the case of SVM results are reported for the RBF kernel, except for *Has risk factor* that reports results for the linear kernel.

